# Dynamic modulation of social gaze by sex and familiarity in marmoset dyads

**DOI:** 10.1101/2024.02.16.580693

**Authors:** Feng Xing, Alec G. Sheffield, Monika P. Jadi, Steve W. C. Chang, Anirvan S. Nandy

## Abstract

Social communication relies on the ability to perceive and interpret the direction of others’ attention, and is commonly conveyed through head orientation and gaze direction in humans and nonhuman primates. However, traditional social gaze experiments in nonhuman primates require restraining head movements, significantly limiting their natural behavioral repertoire. Here, we developed a novel framework for accurately tracking facial features and three-dimensional head gaze directions of multiple freely moving common marmosets (*Callithrix jacchus*). By combining deep learning-based computer vision tools with triangulation algorithms, we were able to track the facial features of marmoset dyads within an arena. This method effectively generates dynamic 3D geometrical facial frames while overcoming common challenges like occlusion. To detect the head gaze direction, we constructed a virtual cone, oriented perpendicular to the facial frame. Using this pipeline, we quantified different types of interactive social gaze events, including partner-directed gaze and joint gaze to a shared spatial location. We observed clear effects of sex and familiarity on both interpersonal distance and gaze dynamics in marmoset dyads. Unfamiliar pairs exhibited more stereotyped patterns of arena occupancy, more sustained levels of social gaze across social distance, and increased social gaze monitoring. On the other hand, familiar pairs exhibited higher levels of joint gazes. Moreover, males displayed significantly elevated levels of gazes toward females’ faces and the surrounding regions, irrespective of familiarity. Our study reveals the importance of two key social factors in driving the gaze behaviors of a prosocial primate species and lays the groundwork for a rigorous quantification of primate behaviors in naturalistic settings.

## Introduction

Primates, including humans, exhibit complex social structures and engage in rich interactions with members of their species, which are crucial for their survival and development. Among social stimuli, the face holds paramount importance with specialized neural systems (Deen et al., 2023; Hesse & Tsao, 2020) and is attentively prioritized by primates during much of their social interaction. Notably, the eyes garner the most attention among all facial features, playing a pivotal role in indicating the direction of others’ attention and also possibly their intention (Dal Monte et al., 2015; Emery, 2000; Itier et al., 2007). Indeed, understanding and interpreting the gaze of fellow individuals is a fundamental attribute of the theory of mind (ToM) (Martin & Santos, 2016; Saxe & Kanwisher, 2003). While current studies of social gaze using either realistic stimuli or pairs of rhesus macaques (*Macaca mulatta*) in a controlled laboratory setting (Dal Monte et al., 2016; Mosher et al., 2014; Ramezanpour & Thier 2020; Shepherd et al., 2006; Shepherd & Freiwald, 2018) provide valuable insights into social gaze behaviors, they are nevertheless limited in their ecological relevance. Many laboratory paradigms rely on head-restrained animals and simplified stimuli, decoupling eye movements from natural head–body dynamics and limiting spontaneous social behavior (Land, 2006; Einhauser et al., 2007; Foulsham et al., 2011). Social gaze is also typically examined in tightly controlled dyadic or screen-based paradigms, which lack the reciprocal, context-dependent, and multi-agent dynamics of natural social interactions (Shepherd, 2010; Birmingham et al., 2008). In contrast, gaze behavior in natural settings is shaped by locomotion, body posture, spatial relationships, and social structure—factors largely absent from laboratory studies (Emery, 2000; Klein et al., 2009). Consequently, the generalizability of findings from controlled experiments to freely moving, socially embedded interactions remains unclear, motivating the need for more naturalistic yet quantitative approaches.

To address these limitations, we turned to common marmosets (*Callithrix jacchus*), a highly prosocial primate species known for their social behavioral and cognitive similarities to humans (Miller et al., 2016). Marmosets are also a model system with increasing applications in computational ethology (Mitchell et al., 2014; Ngo et al., 2022). Like humans, they engage in cooperative breeding, a social system in which individuals care for offspring other than their own, usually at the expense of their reproduction (French, 1997; Solomon & French, 1997). Gaze directions, inferred from head orientation, hold crucial information about marmoset social interactions (Heiney & Blazquez, 2011; Spadacenta et al., 2022). The emergence of computational ethology (Anderson & Perona, 2014; Datta, Anderson, Branson, Perona, & Leifer, 2019) has propelled the development of a host of computer vision tools using deep neural networks (e.g., OpenPose by Cao et al., 2017, DeepLabCut by Mathis et al., 2018, DANNCE by Dunn et al., 2021; SLEAP by Pereira et al., 2022). Moreover, we have recently developed a scalable marmoset apparatus for automated pulling (MarmoAAP) for studying cooperation in marmoset dyads (Meisner et al., 2024). However, tracking head gaze direction in multiple freely moving marmosets poses a challenging problem, not yet solved by existing computer vision tools. This problem is further complicated by the fact that accurately tracking gaze directions in primates requires three-dimensional information.

Here, we propose a novel framework based on a modified DeepLabCut pipeline, capable of accurately detecting body parts of multiple marmosets in 2D space and triangulating them in 3D space. By reconstructing the face frame with six facial points in 3D space, we can infer the head gaze of each animal across time. Importantly, marmosets that are not head-restrained use rapid head movements for reorienting and visual exploration (Pandey et al., 2020), and therefore the head direction serves as an excellent proxy for gaze direction in unrestrained marmosets. With this framework in place, we investigated the gaze behaviors of male-female pairs of freely moving and interacting marmosets to quantify their social gaze dynamics. We investigated these gaze dynamics along the dimensions of sex and familiarity and found several key differences along both of these important social dimensions, including increased partner gaze in males that is modulated by familiarity, increased joint gaze among familiar pairs, and increased gaze monitoring by males. This fully automated tracking system can thus serve as a powerful tool for investigating primate group dynamics in naturalistic environments.

## Results

### Experimental setup and reconstruction of video images in 3D space

The experimental setup consisted of two arenas made with acrylic plates that allowed two marmosets to visually interact with each other while being physically separate (Fig. S1A). Each arena was 60.96 cm long, 30.48 cm wide, and 30.48 cm high. Five sides of the arena, except the bottom, were transparent allowing a clear view of the animal subjects under observation. The bottom side of the arena was perforated with 1-inch diameter holes arranged in a hexagonal pattern to aid the animal’s traction. The arenas were mounted on a rigid frame made of aluminum building blocks, with the smaller sides facing each other, and were separated by a distance of 30.48 cm. A movable opaque divider was placed between the arenas during intermittent breaks to prevent the animals from having visual access to each other (Methods). Two monitors were attached to the aluminum frame, one on each end, for displaying video or image stimuli to the animals. To capture the whole experimental setup, two sets of four GoPro 8 cameras were attached to the frame, where each set of cameras captured the view of one of the arenas.

After obtaining the intrinsic parameters of the cameras by calibration and the extrinsic parameters of the cameras by L-frame analysis (see Methods), we established a world coordinate system of each arena surrounded by the corresponding set of four cameras. Crucially, the two independent world coordinate systems of the two arenas were combined by measuring the distance between the two L-shaped frames and adding this offset to one of the world coordinate systems.

With the established world coordinate system, any point captured by two or more cameras could be triangulated into a common three-dimensional space. Thus, the experimental setup was reconstructed into three-dimensional space by manually labeling the vertices of the arenas and monitors in the image space captured by the cameras (Fig. S1B). The cameras on the monitor ends (marked as ‘ML1’, ‘ML2’, ‘MR1’, ‘MR2’ in Fig. S1B) recorded both animal subjects, whereas the cameras in the middle (marked as ‘OL1’, ‘OL2’, ‘OR1’, ‘OR2’ in Fig. S1B) recorded only one animal subject.

### Automatic detection of facial features of two marmosets

Six facial features – the two tufts, the central blaze, two eyes, and the mouth (Fig. 1A) – were selected for automated tracking using a modified version of a deep neural network (DeepLabCut, Mathis et al., 2018). The raw video from each camera was fed into the network to compute the probability heatmaps of each facial feature. We modified the original method to detect features of two animals (Fig. 1B). After processing the raw video, two locations with the highest probability over a threshold (95%) were picked from each probability heatmap (Fig. 1B, *feature detection*). Since all the features from the same animal should be clustered in image space, a K-means clustering algorithm (Fig. 1B, *initial clustering*) was used on the candidate features with the constraint that one animal can only have one unique feature (Fig. 1B, *refine clustering*); for example, one animal cannot have two left eyes. After clustering, two clusters of features corresponding to the two animals were obtained. To detect outliers that were not valid features, we first calculated a distribution of within-cluster distances (Fig. 1B, *remove outliers*). Outliers were determined as those points that had nearest-neighbor distances which were two standard deviations above the average within-cluster distance and were excluded from subsequent analyses. Note that the above analyses were performed independently for each video frame.

**Figure 1.**
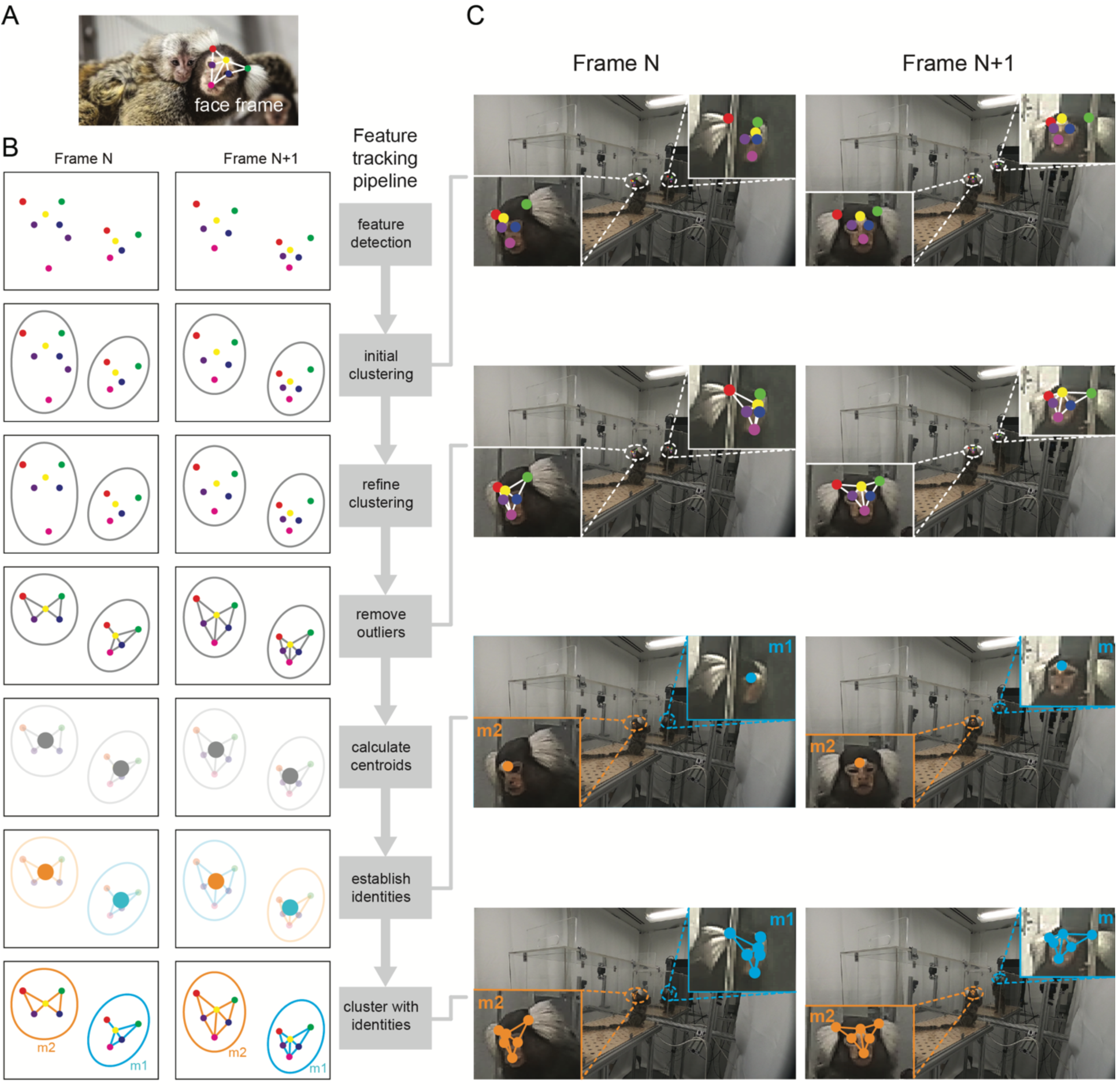
Pipeline of detecting facial features of two marmosets. (A) Six facial features of the marmoset (face frame) are color-coded: right tuft (red), central blaze (yellow), left tuft (green), right eye (purple), left eye (blue), and mouth (magenta). (B) Feature tracking pipeline (right) with the corresponding illustration for each step across two adjacent video frames (left). At the end of the pipeline, the facial features are clustered with the identities assigned consistently across frames. Facial points are color-cod­ed as in A. (C) Example frames of four steps in the pipeline shown in B. It can be seen clearly that the facial points are tracked and clustered accurately, and the identities are consistent across frames.

To establish temporal continuity across video frames and track animal identities, we first calculated the centroid of each cluster (Fig. 1B, *calculate centroids*). Under the heuristic that centroid trajectories corresponding to each individual animal are smoothly continuous in space and time (i.e., there are no sudden jumps or reversals in centroid location across frames at our sampling rate of 30Hz), we assigned identities to the centroids thus enabling us to track identities over time (Fig. 1B, *establish identities*). The facial features corresponding to a centroid inherited this identity (Fig. 1B, *cluster with identities*). We tested our automated pipeline against a manually labeled data set and found that features were detected with high precision (root mean squared error = 3.31 ± 0.1 pixels [mean +/- s.e.m.]; *n* = 2400 facial features; see Methods).

Our method thus allowed us to accurately detect and track the facial features of two marmosets in the arena (Fig. 1C), and in more general contexts such as in a home cage with occlusions (Supplementary Video 1).

### Inferring head-gaze direction in 3D space

In our experimental setup, the cameras in the middle unambiguously recorded only one animal. The centroids of the facial features in the image space recorded by the middle cameras were triangulated into three-dimensional space and were constrained to be confined within the bounds of the arena. Any missing centroids were filled by interpolation from neighboring frames from both the past and future time points. The triangulated centroids acquired from cameras in the middle were then projected into the image space of cameras on the monitor ends. Since both animals were recorded by the cameras on the monitor ends, facial features from these camera views were detected with identities assigned (as described in the 2D pipeline). In the image space of cameras on the monitor ends, the cluster of facial feature points closer to the projected centroid and within the bounds of the arena were kept for later triangulation.

The detected facial features captured by all four cameras for one animal were subjected to triangulation. For each feature, results from all possible pairs of four cameras were triangulated. All the triangulation results were averaged to yield the final coordinates of the body part in three-dimensional space. Any missing features were filled by interpolation from neighboring frames including the previous and future time points. Testing against a manually labeled and triangulated data set yielded a high precision of 3D feature detection (root mean squared error = 3.7 ± 0.1 mm [mean +/- s.e.m.]; *n* = 2400 facial features; see Methods).

The six facial points constituted a semi-rigid geometric frame as the animal moves in 3D space (‘face frame’; Fig. 1), allowing us to infer the animal’s head-gaze direction as follows. The gaze direction was calculated as the normal to the facial plane (‘face norm vector’) defined by the two eyes and the central blaze. The position of the ear tufts, which were behind this facial plane, was used to determine gaze direction. Since marmoset saccade amplitudes are largely restricted to 10 degrees (median less than 4 degrees) (Mitchell et al., 2014), we modeled the head gaze as a virtual cone with a solid angle of 10 degrees (‘gaze cone’) emanating from the facial plane (Fig. 2A). Notably, with multiple camera views, the face frame can be reconstructed even when the face was invisible to one of the cameras, such that the reconstructed face frame in 3D can be projected back into the image space to validate the accuracy of the detection and reconstruction (Fig. 2B). We were thus able to obtain the animal’s continuous movement trajectory and the corresponding gaze direction over time (Fig. 2C, Supplementary Video 2). To characterize stable gaze epochs for subsequent analyses, we calculated the velocity of the face norm vector and applied a stringent threshold of 0.05 (normalized units) below which the marmoset’s head movement was considered stationary (Fig 2D).

**Figure 2.**
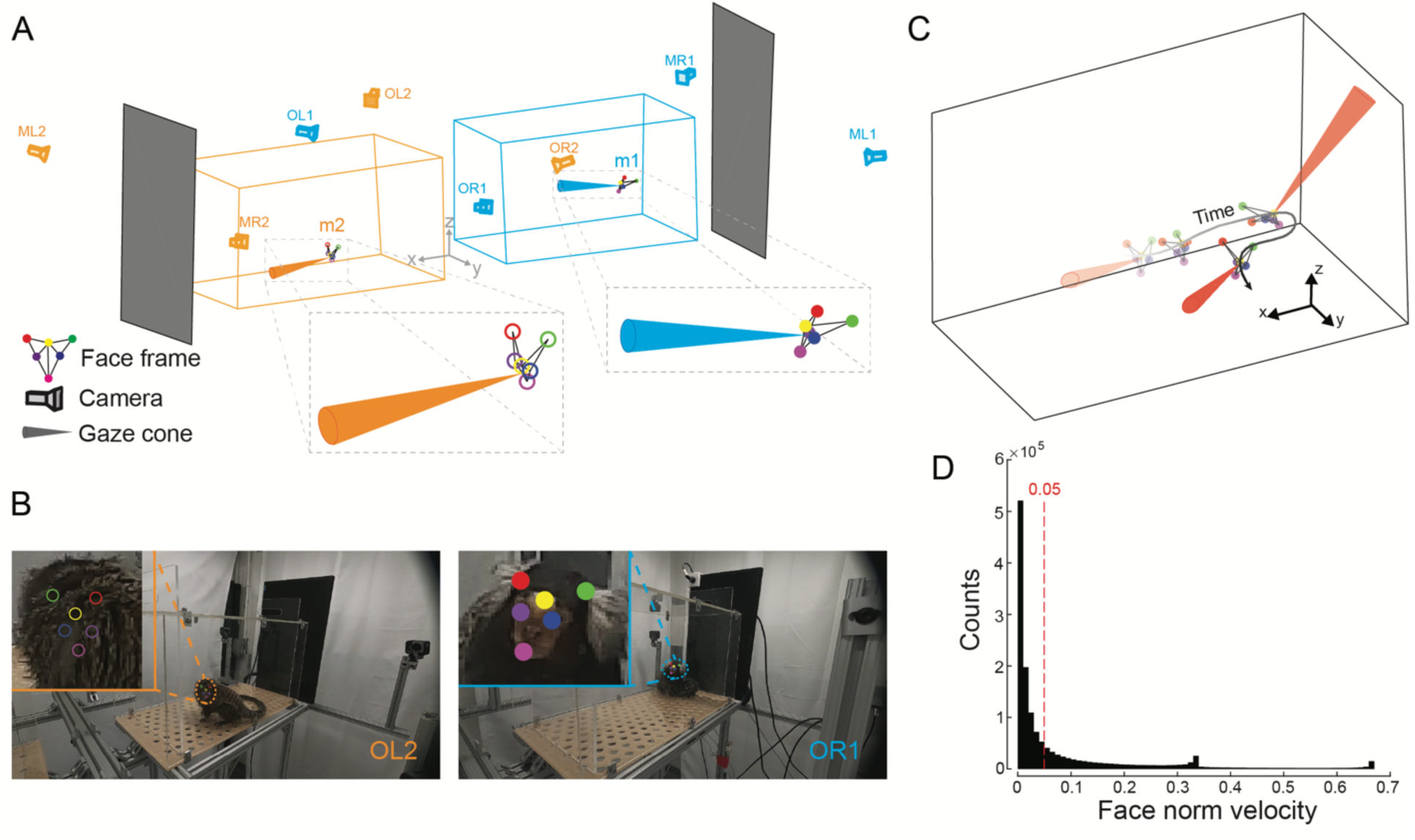
3D Reconstruction of facial features and head gaze modeling. (A) The face frames of two marmosets are reconstructed in 3D using the tracked facial points in Fig. 1A. A cone perpendicular to the face frame (gaze cone; 10-degree solid angle) is modeled as the head gaze. (B) Two example frames with the facial points projected from 3D space onto different camera views are shown. The left frame demonstrates that the facial points can be detected using information from other cameras, even if the face is invisible from that viewpoint. (C) Trajectory of the reconstructed face frame and the corresponding gaze cones across time. (D) The histogram of the face norm velocity. The red dotted line (0.05) is the threshold below which the marmoset head direction is considered to be stationary

We first examined our method’s ability to characterize gaze behaviors of freely moving marmosets by presenting either video or image stimuli to individual animals on a monitor screen (see Methods). We analyzed gaze behavior following the onset of video stimuli presented at different monitor locations (Fig. S2A). The analysis was restricted to video clips in which human annotators confirmed that the marmosets were looking at the monitor. Consistent with prior work in head-fixed marmosets (Mitchell et al., 2014), gaze–monitor intersection centers clustered around the corresponding stimulus locations after stimulus onset, supporting the spatial accuracy of the estimated gaze directions. Marmosets exhibited longer gaze duration to video stimuli compared to image stimuli (Fig. S2B; Mann Whitney U Test, p < 0.001). However, this difference was not caused by differences in gaze dispersion (Fig. S2C; Mann Whitney U Test, ns). By examining the frequency of gaze events, we found that marmosets gazed at the monitor more during the early period of video stimuli presentations compared to the late period (Fig. S2D; Mann Whitney U Test, p < 0.001), while there was no such difference for the image stimuli (Fig. S2D; Mann Whitney U Test, p = 0.4345). There was also a significant difference in gaze frequency between the early period of video presentations compared to the same period for image presentations (Fig. S2D; Mann Whitney U Test, p < 10^-10^). Taken together, our results support that dynamic visual stimuli elicit greater overt attention compared to static stimuli in marmosets, similar to macaques (Dal Monte et al., 2016; Furl et al., 2012) and humans (Chevallier et al., 2015).

### Positional dynamics of marmoset dyads

With this automated system in place, we recorded the behavior of four pairs of familiar marmosets and four pairs of unfamiliar marmosets. We first observed that preferred spatial position was highly non-uniform, with animals preferring to occupy the ends of the elongated arena rather than the center. Data from each pair was recorded in one session consisting of ten five-minute free-viewing blocks interleaved with five-minute breaks. Each pair consisted of a male and a female animal. The familiar pairs were defined as cage mates, while each member of an unfamiliar pair was from a different home cage with no visual access to each other while in the colony. We first examined the movement trajectories of marmoset dyads and used the centroids of the face frames across time to represent the trajectories. For an example 5-minute segment (Fig. 3A), we observed that the marmosets preferred to stay at the two ends of their respective arenas. This was confirmed by a heatmap of projections of the trajectories on the plane parallel to the vertical long side of the arenas (‘XZ’ plane). Furthermore, there were two hotspots along the vertical axis in the heatmap of projections to the vertical short side plane (‘YZ’ plane), suggesting that the animals’ preferred body postures were either upright or crouched.

**Figure 3.**
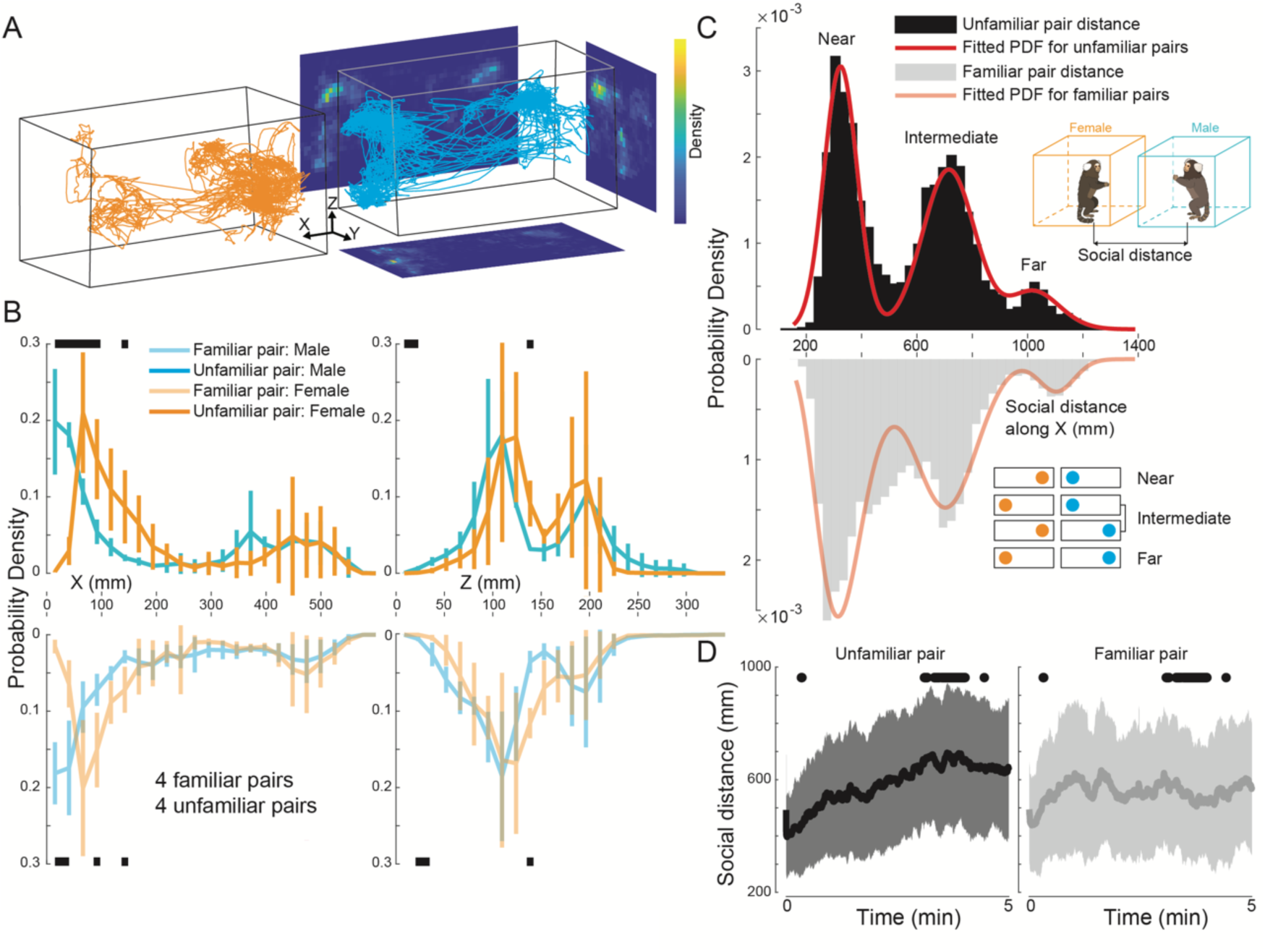
Positional dynamics of marmoset dyads. (A) Movement trajectories of the face frame centroids for a marmoset pair (orange for female, cyan for male) in an example five-minute block. The heatmaps were calculated using the projections of the trajectories to XY, YZ, and XZ planes. (B) Marginal distributions of movement trajectories along the X and Z axes were calculated for all marmosets and grouped by familiarity and sex (transparent colors for familiar pairs, opaque colors for unfamiliar pairs). Black bars indicate significant differences between pairs of distributions (Mann-Whitney U test, significance level at 5%). (C) Histograms of social distance along the X axis show trimodal distributions for both familiar pairs (gray) and unfamiliar pairs (black). The top inset shows a schematic of the arena configuration with corresponding colors (orange for females and cyan for males). Social distance is defined as the Euclidean distance between the two marmosets. The fitted red curve for the unfamiliar pairs is a tri-Gaussian distribution, while the fitted red curve for the familiar pairs is a mixture of Gamma and Gaussian distributions, with the first peak as the Gamma distribution. The three regions were designated as ‘Near’, Intermediate’, and ‘Far’. The bottom inset illustrates the reason for this nomenclature. (D) Temporal evolution of social distance for unfamiliar and familiar pairs (which each 5-minute viewing block). The central dark line is the mean and the shaded area is the standard deviation. Black dots indicate significant differences (Mann-Whitney U test, significance level at 5%).

Second, we found a distinct sexual dimorphism in spatial positioning, where males consistently stayed closer to their partners than females. We examined the marginal distributions of the movement trajectories along the horizontal (‘X’) and vertical (‘Z’) axes across all sessions and grouped them along the dimensions of sex and familiarity (Fig. 3B). Along the X axis (Fig. 3B, left), the distributions were slightly bimodal, with the main peak in the region near to the inner edge of the arenas. Regardless of familiarity, male marmosets tended to stay closer to the inner edge compared to females, as shown by the significant differences in the distributions when X ranged from 0 to 150 mm. However, there were no significant differences between the same sex members of familiar and unfamiliar pairs. For the Z axis (Fig. 3B, right), the distributions were bimodal (Bimodality coefficients (BC) all exceeded the threshold at 5/9 with a 5% margin; BC(familiar male) = 0.5863; BC(familiar female) = 0.6278; BC(unfamiliar male) = 0.6235; BC(unfamiliar female) = 0.6570; Warren Sarle’s bimodality test), consistent with what we observed in the heatmaps indicating either upright or crouched postures. The positional distributions along the Z axis were not different based on sex or familiarity.

Third, our analysis of dyadic distance revealed that familiar pairs spend more time in close proximity and maintain more flexible social distances over time compared to unfamiliar pairs. To characterize the social distance dynamics of the freely moving dyads, we calculated the distance between the centroids of the pairs. We then examined the distributions of the social distance along the X-axis separately for familiar and unfamiliar pairs (Fig. 3C). The distributions were trimodal, and can be explained by the bimodal distribution of movement trajectories of individual marmosets along the X-axis. As mentioned above, marmosets tended to stay at the two ends of their arenas, and thus, combinations of preferred positions at the two ends for the dyads (see insets in Fig. 3C) resulted in the trimodal distribution. We termed these three peaks as ‘Near’, ‘Intermediate’, and ‘Far’. To quantify these distributions, we fitted the empirical data with mixture models using maximum likelihood estimation (see Methods). The social distance for unfamiliar pairs was best fitted by a mixture of three Gaussians while the distribution for familiar pairs was best fitted by a mixture of Gamma and Gaussian distributions. The first peak (‘Near’) of the familiar-pair distribution was best fitted by a Gamma distribution, implying a higher degree of dispersion when familiar marmosets were close to each other. Upon examining the temporal evolution of the social distance (within each 5-minute viewing block), we found that the social distance of unfamiliar pairs increased over time, whereas this distance fluctuated over time for familiar pairs (Fig. 3D), further indicating that the social distance dynamics of marmoset dyads depended on familiarity.

### Social gaze dynamics of marmoset dyads

We next investigated the interactive aspects of gaze behaviors in freely moving marmoset dyads. We found that social interaction in marmosets is characterized by a distinct sexual dimorphism in gaze interest and a transition from “monitoring” in unfamiliar pairs to “active reciprocation” in familiar pairs. The gaze interaction between two animals could be simplified as the relative positions of two gaze cones in three-dimensional space (see Methods; Fig. 4A; Supplementary Video 3). If the gaze cone of one animal intersected with the facial plane of the second animal (but not vice versa), we termed it ‘partner gaze’. If the gaze cones of both animals intersected with that of the other’s facial place, we termed it ‘reciprocal gaze’. In our dataset, the instances of reciprocal gaze were very low and were thus excluded from further analysis. If the two cones intersected anywhere outside the facial planes, we termed it ‘joint gaze’. All other cases were regarded as ‘no interaction’ between the two animals.

**Figure 4.**
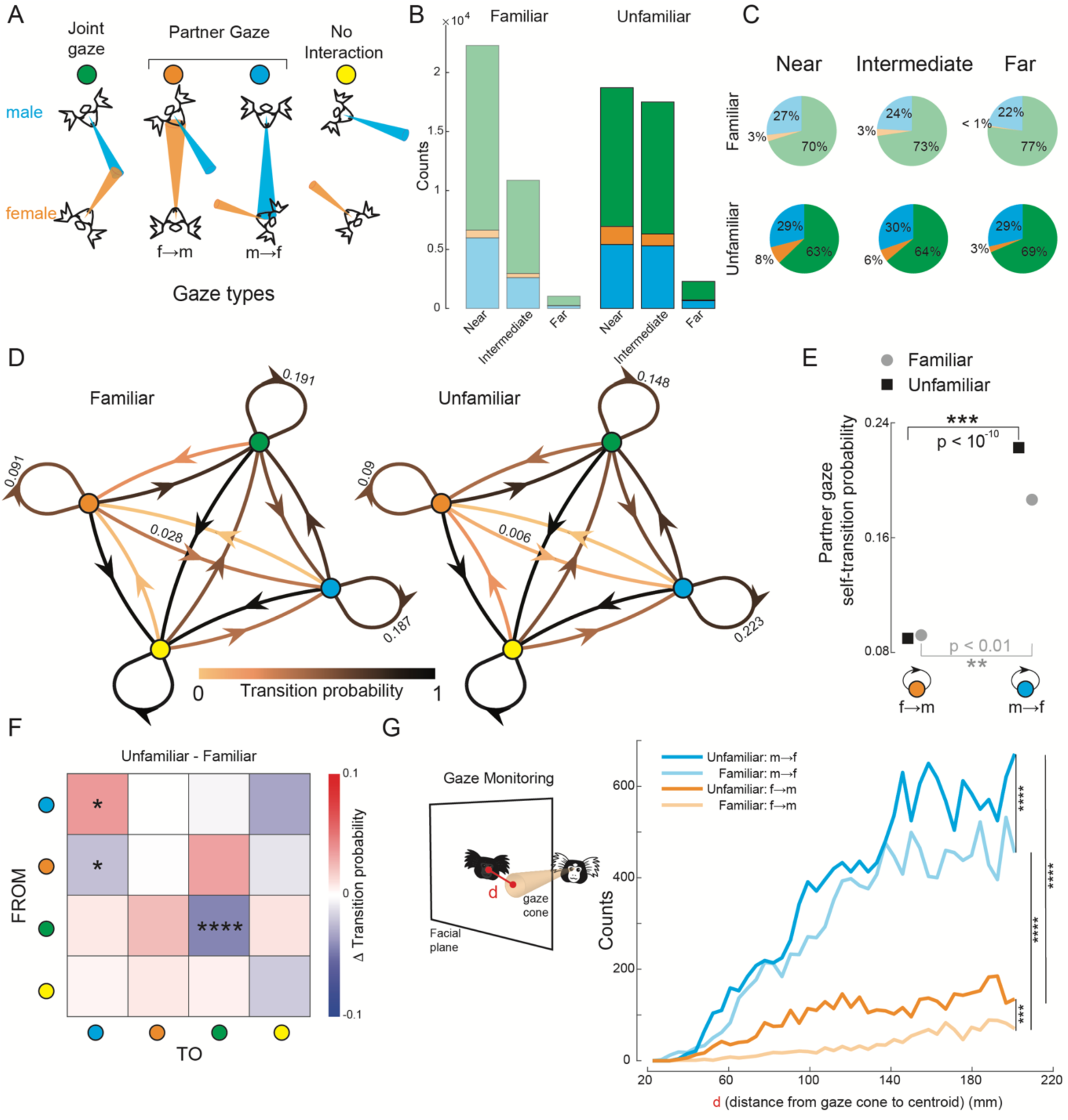
Live interactive gaze analysis of unfamiliar and familiar marmoset dyads. (A) Gaze type categorized based on the relative positions of the gaze cones. Joint gaze is defined as two marmosets looking at the same location. A partner gaze is defined as one animal looking at the other animal’s face (but not vice versa). No interaction occurs when the two gaze cones do not intersect. (B) Histograms of gaze count as a function of inter-animal distance, shown separately for familiar and unfamiliar pairs. (C) Same data as in B shown as pie charts of percentages in social gaze states. (D) Gaze state transition diagrams for familiar and unfamiliar pairs. The nodes are the gaze states and the edges connecting the nodes represent the transition between states. Edge colors indicate transition probabilities. (E) Partner gaze self-transition probabilities for familiar and unfamiliar pairs (χ2 test). (F) Delta transition matrix between the unfamiliar pair and familiar pair state transition diagrams. Transitions that are significantly different across familiarity are marked by asterisks (χ2 test, male to male, p < 0.05; female to male, p < 0.05; joint to joint, p < 0.0001). (G) Left, The schematic illustrates how gazing toward the surrounding region of a partner’s face area was measured. Right, Counts of gaze towards the surrounding region of the partner’s face by familiarity and sex. (Mann-Whitney U test, “* means p < 0.001; **‘* means p < 0.0001)

Our analysis of gaze states reveals that male marmosets maintain a high baseline of interest in females, though this relationship depends on familiarity and spatial proximity. We identified stable gaze epochs by thresholding the face norm vector velocity (Fig. 2D). Stable epochs were categorized into gaze states based on the gaze event types described above. We first analyzed the fraction of gaze states in the three position ranges identified from the social distance analysis (Fig. 4B,C). We found that male marmosets gazed more toward their partner females’ faces regardless of familiarity (p < 0.01, *χ*^2^ test). However, while male interest in familiar females decreased as social distance increased, interest in unfamiliar females remained constant across different social distances, suggesting a persistent monitoring of novel partners (Fig. 4C). Moreover, females in unfamiliar pairs exhibited significantly higher partner gazes (female→male) compared to those in familiar pairs (p < 0.01, *χ*^2^ test; Fig. 4C). The total counts of social gaze states (joint gaze and partner gaze) were higher for familiar pairs when they were near, but these decreased more dramatically with increasing distance (Fig. 4B).

Our state-transition analysis revealed that gaze dynamics are governed by a combination of persistent male attention and a shared tendency for males to follow the female’s gaze into joint attention. To investigate the dynamics of these gaze states, we computed state transition probabilities among distinct gaze event types for familiar and unfamiliar dyads. Markov chain modeling of state transitions further highlighted that familiar dyads engage in more complex, reciprocal social exchanges compared to the one-sided monitoring seen in unfamiliar dyads (Fig. 4D). We first focused on the recurrent (self-transition) edges for the partner gaze states. Recurrent edges indicate a transition back to the same stable gaze state after a break likely due to physical movement, and reflect the robustness of the state despite movement. In line with our previous results (Fig. 4B,C), males exhibit significantly higher recurrent partner gazes compared to females, irrespective of familiarity (*χ*^2^ test, unfamiliar male vs unfamiliar female, p < 10-10; familiar male vs familiar female, p < 0.01; Fig. 4E). Furthermore, we observed a consistent pattern of gaze-following across both groups; approximately 17% of male-to-female partner gazes transitioned directly into joint gaze (Fig. 4D), suggesting that males frequently utilize the female’s gaze orientation to guide their own environmental exploration.

Our comparison of state transition probabilities reveals that familiarity transforms male gaze behavior from a persistent monitoring of novel partners to a more socially responsive, reciprocal interaction (Fig. 4F). First, recurrent male partner gaze (male→female) was significantly enhanced in unfamiliar pairs (p < 0.05, *χ*^2^ test), suggesting a heightened interest in unfamiliar females. Second, there was a higher probability of transition from a female partner gaze to a male partner gaze in familiar pairs compared to unfamiliar pairs, suggesting that familiar males have a greater awareness of and tendency to reciprocate their partners’ gaze (p < 0.05, *χ*^2^ test). Third, there was a higher probability of recurrent joint gazes in familiar pairs compared to unfamiliar pairs, suggesting that familiar pairs explore common objects more than unfamiliar pairs (p < 10-4, *χ*^2^ test).

Monitoring others to anticipate their future actions is critical for successful social interactions (Hari et al., 2015). In particular, successful interactive gaze exchanges require constant monitoring of others’ gaze. In addition to direct eye contact, we found that unfamiliar marmosets engage in increased peripheral monitoring of their partners. We analyzed the gaze distribution in the surrounding region of a partner’s face to estimate gaze monitoring tied to increased social attention (Dal Monte et al., 2022). We quantified this by the distance between the centroid of the partner’s face-frame and the point of intersection of the gaze cone with the partner’s facial plane (Fig. 4G, left). Unfamiliar marmosets (both males and females) showed significantly higher (Mann-Whitney U test, unfamiliar male vs familiar male, p < 0.0001; unfamiliar female vs familiar female, p < 0.001) incidences of gaze toward the surrounding region of the partner’s face (Fig 4G, right; compare darker lines with the lighter lines). Further, males exhibited markedly higher (Mann-Whitney U test, p < 0.0001) incidences of gaze toward the partner females’ face (Fig 4G, right; compare cyan lines with the orange lines).

Finally, our analysis of joint gaze distribution underscores that familiar pairs establish shared attention more flexibly across their social distance, whereas shared attention in unfamiliar pairs are constrained by spatial proximity. Joint gaze is crucial in primates as it underpins social communication and coordination, serving as a foundation for more complex behaviors like cooperation and shared attention (Emery, 2000; Tomasello et al., 2005). We analyzed joint gaze behaviors between marmoset dyads, first projecting the locations of joint gazes onto different 2D planes around the arena (Fig. 5A). Significant asymmetry was observed in the distribution of joint gazes on the ‘XY’ and ‘XZ’ planes, with the majority of joint gazes occurring within the female’s arena, regardless of familiarity. This is consistent with our earlier results showing more partner-directed gazes from males and increased attention to the region surrounding the female partner (Fig. 4B,C,G). Notably, joint gazes between familiar pairs were more centrally distributed compared to those between unfamiliar pairs. When examining the ‘XZ’ and ‘YZ’ planes, we found that most joint gazes were concentrated along the lower Z-axis, indicating that marmoset pairs tend to establish shared attention closer to ground level. Given the importance of social distance in shaping marmoset behaviors, we further investigated the distribution of joint gazes as a function of social distance. Striking differences were found between familiar and unfamiliar pairs (Fig. 5B). The distribution for familiar pairs was best fitted by a tri- Gaussian model, consistent with the distribution of social distances observed in Fig. 3C, suggesting that familiar pairs can establish joint gazes regardless of distance. In contrast, the distribution for unfamiliar pairs was best described by a lognormal model, indicating that unfamiliar pairs tend to establish joint gazes only when in close proximity.

**Figure 5.**
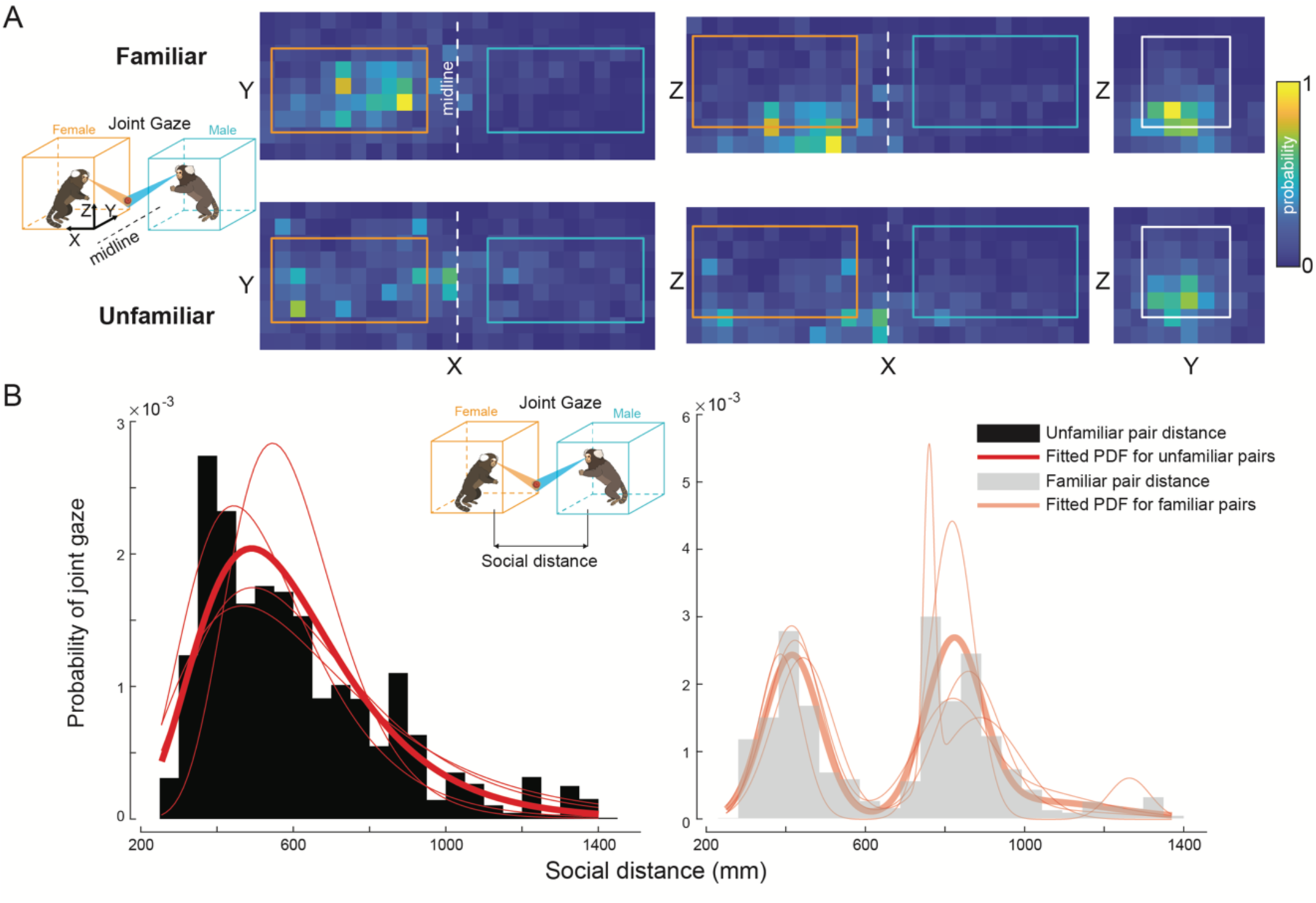
Joint gaze analysis between familiar and unfamiliar dyads. (A) Heatmaps of joint gaze locations projected onto 2D planes (From left to right: ‘XY’, ‘XZ’, ‘YZ’). The icon on the left show a schematic of a joint gaze along with the arena configuration with corresponding colors (orange for females and cyan for males). The colored rectangles superimposed on the heatmaps show the projections of the arenas onto those 2D planes. The white dotted line indicates the midline between the two arenas. (B) The probability of joint gaze at varying social distances shows distinct distributions for unfamiliar pairs (black) and familiar pairs (gray). The fitted red curve for the unfamiliar pairs is a lognormal distribution, while the fitted red curve for the familiar pairs is a tri-Gaussian distribution. Thin fitted red curves in both panels are the data from individual pairs. The inset shows a schematic of a joint gaze and the social distance metric. Social distance is defined as the Euclidean distance between the two marmosets.

Overall, we found that both the social dimensions we examined – familiarity and sex – are significant determinants of natural gaze dynamics among marmosets.

## Discussion

In this study, we first presented a novel framework for the automated, markerless, and identity-preserving tracking of 3D facial features of multiple marmosets. By building on top of the deep-learning framework provided by DeepLabCut, we used a constellation of cameras to overcome “blindspots” due to occlusion and imposed spatiotemporal smoothness constraints on the detected features to establish and preserve identities across time. The tracked facial features from each animal form a semi-rigid face frame as the animal moves freely in 3D space, thereby allowing us to infer the animal’s gaze direction at each moment in time. It is important to reiterate that unrestrained marmosets use rapid saccadic head-movements for reorienting (Pandey et al., 2020) and have a limited amplitude range of saccadic eye-movements (Mitchell et al., 2014). Thus their head direction serves as excellent proxy for gaze direction in unrestrained conditions.

Current methods for tracking and estimating animal body postures, such as DeepLabCut (Mathis et al., 2018) and SLEAP (Pereira et al., 2022), offer substantial advantages in precision and flexibility, particularly through the use of deep learning for markerless tracking across diverse species. These methods excel in laboratory settings, where they can achieve high accuracy without invasive markers and are adaptable for various animals and tasks. However, many of these tools struggle with occlusion, especially in crowded environments or natural habitats, limiting their effectiveness in complex social interactions or open-field studies (Sturman et al., 2020). Additionally, when tracking marmosets that exhibit significant vertical movement, requiring 3D tracking, the performance of these methods is suboptimal. While 3D tracking systems like DANNCE (Dunn et al., 2021) offer enhanced spatial accuracy, these methods are limited to single-animal tracking, which is insufficient for studies in social neuroscience.

Primates are highly visual species whose physical explorations of their environment are not confined to two-dimensional surfaces. Gaze is a critical component of primate social behavior and conveys important social signals such as interest, attention, and emotion (Emery, 2000). Assessment of gaze is therefore important to understand non-verbal communication and interpersonal dynamics. Our 3D gaze tracking approach was able to capture both the positional and gaze dynamics of freely moving marmoset dyads in a naturalistic context. We observed clear effects of sex and familiarity on both interpersonal and gaze dynamics. Unfamiliar pairs exhibited more stereotyped patterns of arena occupancy, more sustained levels of social gaze across distance, and increased gaze monitoring, suggesting elevated levels of social attention and the need to constantly track the movements of a novel conspecific compared to familiar pairs. On the other hand, familiar pairs exhibited more recurrent joint gazes in the shared environment compared to unfamiliar pairs. Critically, familiar males also showed a higher tendency to reciprocate their partner’s gaze, suggesting a greater awareness of their partner’s social gaze state and a transition from mere monitoring to active social engagement.

Supported by the natural ethology of marmosets (Yamamoto et al., 2014; Solomon & French, 1997), we found dramatic sex differences in gaze behaviors, with males exhibiting significantly elevated levels of gaze toward females’ faces and the surrounding regions irrespective of familiarity. It is important to note that dominance in marmosets is not strictly determined by gender, as it can vary based on individual personalities and intra-group social dynamics, although breeding females typically dominate social activity within a group (Digby, 1995; Mustoe et al., 2023). While we have not explicitly controlled for dominance in this study, whether part of the observed differences can be attributed to dominance effects needs further exploration.

The distinction between social monitoring and social coordination is further reflected in the spatial flexibility of joint attention. Gaze following plays a crucial role in social communication for humans and nonhuman primates, allowing for joint attention (Emery et al., 1997; Brooks & Meltzoff, 2005; Burkart & Heschl, 2006; Shepherd, 2010). Previous research demonstrated that head-restrained marmosets exhibited preferential gazing toward marmoset face stimuli observed by a conspecific in a quasi-reflexive manner during a free-choice task (Spadacenta et al., 2019). Here we found that males consistently followed the gaze of females into states of joint attention. This ‘gaze following’ occurred with similar frequency in both familiar and unfamiliar pairs, suggesting that monitoring a partner’s gaze to coordinate shared attention is a fundamental component of marmoset social ethology, regardless of the strength of the social bond. Interestingly, we did not find any differences in gaze-following behaviors (transition from partner gaze to joint gaze) along the social dimensions we tested here. Future investigation of such behaviors by manipulating social variables such as dominance or kinship could provide a comprehensive understanding of gaze following and joint attention in naturalistic behavioral contexts. The scarcity of reciprocal gazes in our study may be attributed to the task-free experimental setup employed. Indeed, in other joint action tasks requiring cooperation for rewards, marmosets actively engage in reciprocal gaze behaviors (Miss & Burkart, 2018).

While we focused on the tracking of facial features in this study, our automated system has the potential to extend to 3D whole-body tracking, encompassing limbs, tail, and the main body features of marmosets. In our system, multiple cameras surrounding the arena ensure that each body part of interest can be tracked through at least two cameras, enabling triangulation in 3D space. Our current system uses a pre-trained ResNet model (He et al., 2015) to track body parts of interest. However, considering the challenges posed by whole-body tracking, such as interference from marmosets’ fur that complicates feature detection, the adoption of cutting-edge transformer networks like the vision transformer model (Dosovitskiy et al., 2020) might significantly improve detection performance. Such an advancement in tracking and reconstructing the entire marmoset body frame would enable the analysis of such data using unsupervised learning techniques (Berman et al., 2014; Calhoun et al., 2019) and thereby provide a deeper understanding of primate social behavior.

In summary, our study lays the groundwork for a rigorous quantification of primate behaviors in naturalistic settings. Not only does this allow us to gain deeper insights beyond what is possible from field notes and observational studies, but it is also a key first step to go beyond current reductionist paradigms and understanding the neural dynamics underlying natural behaviors (Miller et al., 2022).

## Acknowledgments

This research was supported by the National Institute of Mental Health (R21 120672, SWCC, ASN, MPJ), Simons Foundation Autism Research Initiative (SFARI 875855, SWCC, ASN, MPJ), Brain Research Foundation (BRFSG-2020-05, ASN), Yale Orthwein Scholar Funds (ASN) and by the National Eye Institute core grant for vision research (P30 EY026878 to Yale University). We would like to thank the veterinary and husbandry staff at Yale for excellent animal care. We would like to thank Weikang Shi for helpful discussion on the manuscript.

## Author contributions

ASN, SWCC & MPJ conceptualized the project. FX collected the data with assistance from AGS. FX analyzed the data. ASN supervised the project. FX, ASN, SWCC & MPJ wrote the manuscript.

## Declaration of interests

The authors declare no competing interests.

## Inclusion and Ethics

We support inclusive, diverse, and equitable conduct of research.

## Methods

### Camera calibration

All cameras (GoPro 8) were calibrated using an 8-by-9 black-white checkerboard. For each camera, the checkerboard was placed at various locations to sample the space of the camera’s field of view. To achieve better calibration performance, the checkerboard was tilted and rotated to varying degrees thus producing a range of different views (Zhang, 2000). The corners of the checkerboard were automatically detected via a standard algorithm (detectCheckerboardPoints() function in the Image Processing and Computer Vision toolbox in MATLAB). The intrinsic parameters of each camera were estimated based on the data obtained from the checkerboard corner detection algorithm (estimateCameraParameters() function in Image Processing and Computer Vision toolbox in MATLAB).

### L-frame analysis

L-shaped frames were used to obtain the extrinsic parameters of the cameras, the rotation matrix, and the transition vector (Timothy et al., 2021). The L-shaped frame was captured by four cameras that recorded one arena. Four points that were unevenly distributed on the L-shaped frame were manually labeled. The information of transformation from world coordinates to camera coordinates was then extracted based on the labeled result (cameraPoseToExtrinsics() function in Image Processing and Computer Vision toolbox in MATLAB).

### Camera recording

GoPro 8 cameras were used and were simultaneously controlled via a Bluetooth remote control (The Remote by GoPro). Videos were recorded at 30 frames/sec with a linear lens. Frame resolution was set at 1920×1080 pixels. A circular polarizer filter was used to mitigate reflection artifacts.

### Deep convolutional neural network (DCNN) model training

We used a modified version of DeepLabCut (Mathis et al., 2018) to perform automated markerless tracking of body parts of interest from two marmosets. The model was trained on 700 hand-labeled image frames extracted from videos of animals in their colony settings. Each image frame was labeled with six facial points: the two tufts, the central blaze, two eyes, and the mouth. The model was trained using GPUs on a large computing cluster for 250,000 iterations until the loss reached a plateau.

### Method validation in both 2D and 3D spaces

To validate the tracking method in 2D space, we manually labeled 200 continuous frames containing two marmosets from the recorded videos. The model was then provided with these 200 frames to detect the body parts. We calculated the differences between the ground-truth coordinates and the detected body parts using the root mean square error (RMSE):

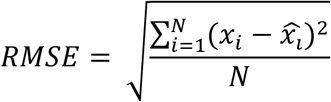

where *x*_*i*_ is the ground-truth coordinate, 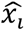 is the detected coordinate, *N* is the total number of body parts.

It is important to note that in cases where certain body parts were occluded in the frames, their coordinates were labeled as (0, 0).

To test the method in 3D space, we manually labeled 200 consecutive pairs of frames containing two marmosets, extracted from two simultaneously recording cameras. The coordinates were triangulated into 3D space. We then fed the model with these 200 pairs of frames to obtain the detected body part coordinates in 3D. The differences between the ground-truth coordinates and the detected body parts were calculated using the same RMSE metric.

### Gaze cone calculation

At each time frame, the gaze direction was calculated as the normal to the facial plane (‘face norm vector’) defined by the two eyes and the central blaze. The position of the ear tufts, which were behind this facial plane, was used to determine the direction of gaze. A gaze cone was defined as a virtual cone of 10-degree solid angle around this norm.

### Head gaze velocity calculation and stable epoch identification

We used the change of the norm over consecutive time frames to calculate the head gaze velocity:

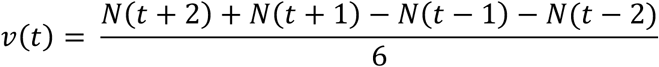

where v(t) is the velocity at time point t, N(t) is the face norm vector at time point t.

We remove all time points where the head gaze velocity was larger than 0.05 in normalized units. Segments no shorter than three consecutive time frames were identified as stable epochs.

### Cone-monitor plane intersection

We modified an existing method (Calinon & Billard, 2006; Sylvain, 2009) to determine the elliptical intersection of a gaze cone and the finite plane defined by the monitor.

### Cone-facial plane intersection

We used a numerical method to determine whether the gaze cone of one animal intersected with the facial plane of the other. The facial plane was defined as the finite triangular plane formed by three facial features: two eyes and mouth. Any point X in 3D within the volume bounded by the cone satisfies the inequality:

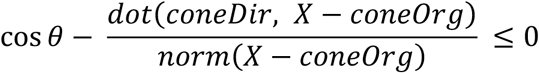

where *θ* is the solid angle of the gaze cone, *coneDir* is the direction vector of the gaze cone, *coneOrg* is the origin point of the gaze cone. The facial plane intersects with the cone if any point within the finite plane satisfies the inequality.

### Cone-cone intersection

To calculate the cone-cone intersection, we used the same numerical method as above. If any point X in 3D simultaneously satisfied the following inequalities:

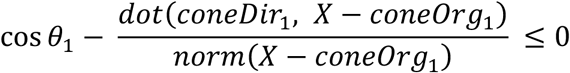

and

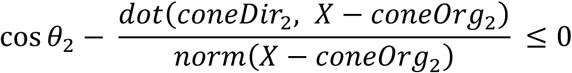

then the two cones were considered to be intersected. Subscripts in the above inequalities indicate the parameters of the two gaze cones under consideration.

### Gaze type definition

To resolve spatial ambiguities where gaze cones might intersect both a partner and an external object, we implemented a hierarchical priority rule. A frame was categorized as Partner Gaze (or Reciprocal Gaze) if the gaze cone intersected the partner’s facial plane, regardless of whether a secondary intersection with the other animal’s gaze cone occurred elsewhere. Joint Gaze was only recorded when both animals’ gaze cones intersected at an external point without either cone containing the partner’s face. This hierarchy ensures that direct social attention is not masked by incidental external intersections.

### Maximum likelihood estimation

We used the maximum likelihood estimation method (mle() function in the Statistics and Machine Learning Toolbox in MATLAB) to fit a mixture of Gamma and Gaussian distributions.

### Markov chain analysis

State transition matrices were obtained based on the behavioral data. These matrices were then used to generate the discrete Markov chains (dtmc() function in Econometrics Toolbox in MATLAB) and plotted (graphplot() function in MATLAB).

### Warren Sarle’s bimodality coefficient

Sarle’s bimodality coefficient is used to test for bimodality. The coefficient is calculated using the MATLAB function written by Hristo Zhivomirov (2024). The code is based on the theory in Pfister et al., 2013.

## Experimental model and subject details

### Animals

Nine adult marmosets were used in this study (four males, five females)[Add info about age ranges of each sex]. Four familiar male/female pairs were each from the same cage. Four unfamiliar male/female pairs were selected from the nine animals such that each member of a pair were from different home cages and did not have visual access to each other while in the colony. Animals were kept in a colony maintained at around 75°F, 60% humidity and a 12h:12h light-dark cycle.

### Single marmoset gazing at the monitor

A single freely moving marmoset was recorded by four cameras surrounding the arena. Video or image stimuli were displayed at one of five locations (Center, Up, Down, Left and Right) on the monitor (location chosen randomly). Each session contained only one stimulus category (either video or image) and consisted of five blocks. Each block consisted of ten five-second stimuli interleaved with ten five-second breaks. Each block started with a white dot in the center of the screen on a black background lasting for one second. At the end of the block, a juice reward (diluted condensed milk, condensed milk : water = 1:7) was delivered with a syringe pump system (NE-500 programmable OEM syringe pump from Pump Systems Inc.) along with an auditory cue.

### Freely interacting marmoset dyads

Two freely moving marmosets, in separate arenas, were recorded by two sets of four cameras surrounding the arenas. Each session consisted of ten five-minute free-viewing blocks interleaved with nine five-minute breaks. A juice reward (diluted condensed milk, condensed milk : water = 1:7) was delivered every minute through two syringe pump systems during the free-viewing blocks. During the breaks, a divider was placed between the two arenas that prevented the marmosets from seeing each other.

## Supplementary Material

**Supplementary Figure 1.**
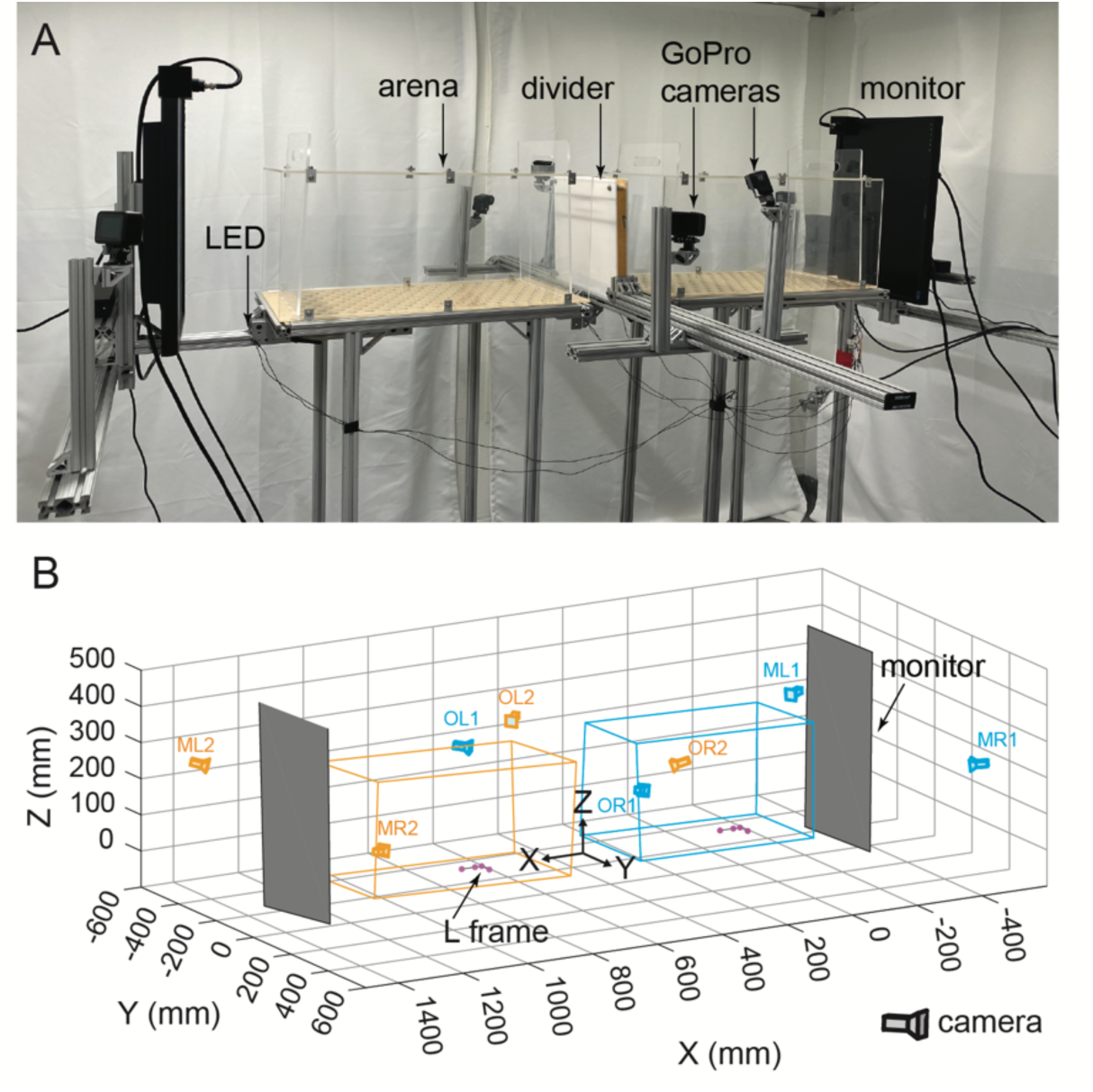
Experimental setup and reconstruction in 3D space. (A) Two transparent acrylic arenas allowed marmosets to visually interact with each other. An opaque divider between the arenas was introduced intermittently to prevent visual access between animals. Twc monitors on two ends were used to display video or image stimuli. Eight cameras surrounding the arenas ensured full coverage of both animals. LEDs at four positions were used to synchronize the video recordings across the set of cameras. (B) 3D reconstruction of the experimental setup. The two arenas are color-coded as orange and cyan. The cameras colored the same as the arenas indicate that they primarily record the marmoset in the corresponding arena. Two purple L-frames within the arenas were used to establish a world coordinate system in the reconstruction process. Two gray planes on both ends are the reconstructed monitors.

**Supplementary Figure 2.**
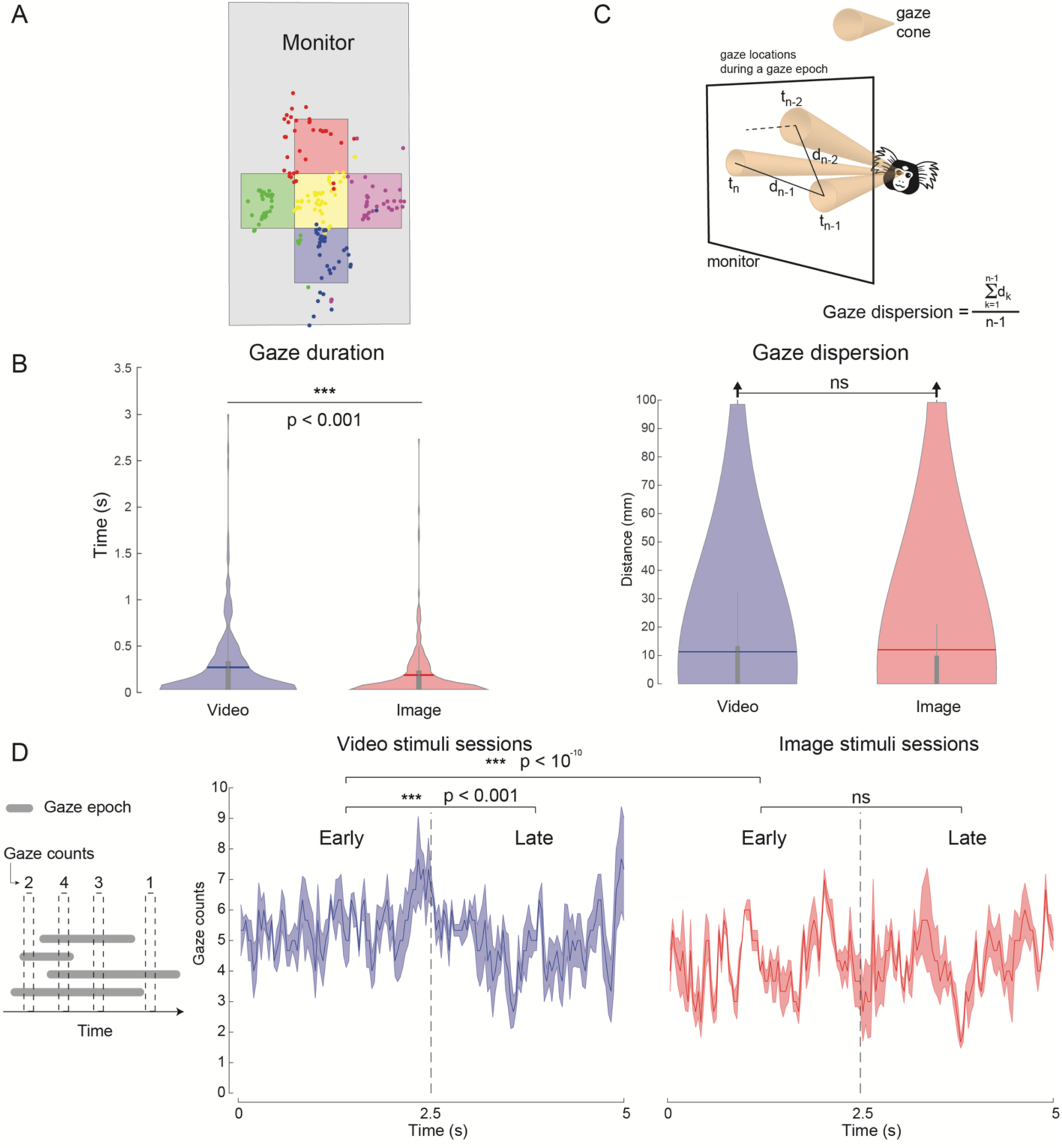
Gaze behavior analysis of a single marmoset viewing stimuli on the monitor. (A) Spatial distribution of gaze-monitor intersection centers during video viewing by a single marmoset. Different colors indicate stimuli presented at different locations on the monitor. Dots of the corresponding colors denote the centers of gaze-cone intersections with the monitor plane one second after stimulus onset. (B) Gaze duration for video stimuli is significantly higher compared to image stimuli (Mann Whitney U Test, p < 0.001). (C) Top, Gaze dispersion is defined as the average distance between the centers of the intersection of the gaze cone and the monitor within a gaze epoch. There was no difference between the video and image stimuli for gaze dispersion (Mann Whitney U Test, ns). (D) Left, Illustration of four gaze epochs (gray bars) to repeated presentations of a stimulus and the gaze counts at different time points within the duration of the presentation. Marmosets have more gazes in the early period than the late period for the video stimuli (Mann Whitney U Test, p < 0.001), however, this is not the case for the image stimuli (Mann Whitney U Test, ns). During the early period, marmosets had significantly higher gaze counts for the video stimuli than the image stimuli (Mann Whitney U Test, p < 10^-10^).

**Video 1**. Results from different processing stages of the facial features detection pipeline are shown for two marmosets in their home cage.

**Video 2**. 3D reconstruction of the face frame and the inferred gaze cone across time for a single marmoset.

**Video 3**. Categorization of gaze behavior epochs of two freely viewing marmosets and transitions between the defined gaze states.

## Notes

### Competing Interest Statement

The authors have declared no competing interest.

### Summary of Updates

Results and Discussion sections updated to address peer review comments; Supp Fig 2 updated

## References

Anderson, David J., & Perona, P. (2014). Toward a Science of Computational Ethology. Neuron, 84(1), 18–31. 10.1016/j.neuron.2014.09.005

Berman, G. J., Choi, D. M., Bialek, W., & Shaevitz, J. W. (2014). Mapping the stereotyped behaviour of freely moving fruit flies. Journal of The Royal Society Interface, 11(99), 20140672.

Birmingham, E., Bischof, W. F., & Kingstone, A. (2008). Social attention and real-world scenes: The roles of action, competition and social content. The Quarterly Journal of Experimental Psychology, 61(7), 986–998. 10.1080/17470210701410375

Brooks, R., & Meltzoff, A. N. (2005). The development of gaze following and its relation to language. Dev Sci, 8(6), 535–543. doi:10.1111/j.1467-7687.2005.00445.x

Burkart, J., & Heschl, A. (2006). Geometrical gaze following in common marmosets (Callithrix jacchus). J Comp Psychol, 120(2), 120–130. doi:10.1037/0735-7036.120.2.120

Calhoun, A. J., Pillow, J. W., & Murthy, M. (2019). Unsupervised identification of the internal states that shape natural behavior. Nature neuroscience, 22(12), 2040–2049. 10.1038/s41593-019-0533-x

Cao, Z., Simon, T., Wei, S.-E., & Sheikh, Y. (2017). Realtime multi-person 2d pose estimation using part affinity fields. Paper presented at the Proceedings of the IEEE Conference on Computer Vision and Pattern Recognition.

Chevallier, C., Parish-Morris, J., McVey, A., Rump, K. M., Sasson, N. J., Herrington, J. D., & Schultz, R. T. (2015). Measuring social attention and motivation in autism spectrum disorder using eye-tracking: Stimulus type matters. Autism Res, 8(5), 620–628. doi:10.1002/aur.1479

Dal Monte, O., Costa, V. D., Noble, P. L., Murray, E. A., & Averbeck, B. B. (2015). Amygdala lesions in rhesus macaques decrease attention to threat. Nature Communications, 6(1), 10161. doi:10.1038/ncomms10161

Dal Monte, O., Fan, S., Fagan, N. A., Chu, C. J., Zhou, M. B., Putnam, P. T., … Chang, S. W. C. (2022). Widespread implementations of interactive social gaze neurons in the primate prefrontal-amygdala networks. Neuron, 110(13), 2183–2197.e2187. doi:10.1016/j.neuron.2022.04.013

Dal Monte, O., Piva, M., Morris, J. A., & Chang, S. W. (2016). Live interaction distinctively shapes social gaze dynamics in rhesus macaques. J Neurophysiol, 116(4), 1626–1643. doi:10.1152/jn.00442.2016

Datta, S. R., Anderson, D. J., Branson, K., Perona, P., & Leifer, A. (2019). Computational Neuroethology: A Call to Action. Neuron, 104(1), 11–24. 10.1016/j.neuron.2019.09.038

Deen, B., Schwiedrzik, C. M., Sliwa, J., & Freiwald, W. A. (2023). Specialized Networks for Social Cognition in the Primate Brain. Annual Review of Neuroscience, 46(1), 381–401. doi:10.1146/annurev-neuro-102522-121410

Digby, L. J. (1995). Social organization in a wild population ofCallithrix jacchus: II. Intragroup social behavior. Primates, 36(3), 361–375. doi:10.1007/BF02382859

Dosovitskiy, A. (2020). An image is worth 16×16 words: Transformers for image recognition at scale. arXiv preprint arXiv:2010.11929.

Dunn, T. W., Marshall, J. D., Severson, K. S., Aldarondo, D. E., Hildebrand, D. G. C., Chettih, S. N., Wang, W. L., Gellis, A. J., Carlson, D. E., Aronov, D., Freiwald, W. A., Wang, F., & Ölveczky, B. P. (2021). Geometric deep learning enables 3D kinematic profiling across species and environments. Nature methods, 18(5), 564–573. 10.1038/s41592-021-01106-6

Einhäuser W, Schumann F, Bardins S, Bartl K, Böning G, Schneider E, König P. Human eye-head co-ordination in natural exploration. Network. 2007 Sep;18(3):267–97. doi: 10.1080/09548980701671094. PMID: 17926195.

Emery, N. J. (2000). The eyes have it: the neuroethology, function and evolution of social gaze. Neurosci Biobehav Rev, 24(6), 581–604. doi:10.1016/s0149-7634(00)00025-7

Emery, N. J., Lorincz, E. N., Perrett, D. I., Oram, M. W., & Baker, C. I. (1997). Gaze following and joint attention in rhesus monkeys (Macaca mulatta). J Comp Psychol, 111(3), 286–293. doi:10.1037/0735-7036.111.3.286

Foulsham T, Walker E, Kingstone A. The where, what and when of gaze allocation in the lab and the natural environment. Vision Res. 2011 Sep 1;51(17):1920–31. doi: 10.1016/j.visres.2011.07.002. Epub 2011 Jul 23. PMID: 21784095.

French, J. A. (1997). Proximate regulation of singular breeding in callitrichid primates. Cooperative breeding in mammals, 34-75.

Furl, N., Hadj-Bouziane, F., Liu, N., Averbeck, B. B., & Ungerleider, L. G. (2012). Dynamic and static facial expressions decoded from motion-sensitive areas in the macaque monkey. J Neurosci, 32(45), 15952–15962. doi:10.1523/jneurosci.1992-12.2012

Hari, R., Henriksson, L., Malinen, S., & Parkkonen, L. (2015). Centrality of Social Interaction in Human Brain Function. Neuron, 88(1), 181–193. 10.1016/j.neuron.2015.09.022

He, K., Zhang, X., Ren, S., & Sun, J. (2016). Deep residual learning for image recognition. In Proceedings of the IEEE conference on computer vision and pattern recognition (pp. 770–778).

Heiney, S. A., & Blazquez, P. M. (2011). Behavioral responses of trained squirrel and rhesus monkeys during oculomotor tasks. Exp Brain Res, 212(3), 409–416. doi:10.1007/s00221-011-2746-4

Hesse, J. K., & Tsao, D. Y. (2020). A new no-report paradigm reveals that face cells encode both consciously perceived and suppressed stimuli. eLife, 9, e58360. doi:10.7554/eLife.58360

Itier, R. J., Alain, C., Sedore, K., & McIntosh, A. R. (2007). Early face processing specificity: it’s in the eyes! J Cogn Neurosci, 19(11), 1815–1826. doi:10.1162/jocn.2007.19.11.1815

Klein, J. T., Shepherd, S. V., & Platt, M. L. (2009). Social attention and the brain. Current biology : CB, 19(20), R958–R962. 10.1016/j.cub.2009.08.010

Land MF. Eye movements and the control of actions in everyday life. Prog Retin Eye Res. 2006 May;25(3):296–324. doi: 10.1016/j.preteyeres.2006.01.002. Epub 2006 Mar 3. PMID: 16516530.

Martin, A., & Santos, L. R. (2016). What Cognitive Representations Support Primate Theory of Mind? Trends Cogn Sci, 20(5), 375–382. doi:10.1016/j.tics.2016.03.005

Mathis, A., Mamidanna, P., Cury, K. M., Abe, T., Murthy, V. N., Mathis, M. W., & Bethge, M. (2018). DeepLabCut: markerless pose estimation of user-defined body parts with deep learning. Nature neuroscience, 21(9), 1281–1289.

Meisner Olivia C, Shi Weikang, Fagan Nicholas A, Greenwood Joel, Jadi Monika P, Nandy Anirvan S, Chang Steve WC (2024) Development of a Marmoset Apparatus for Automated Pulling (MarmoAAP) to Study Cooperative Behaviors eLife 13:RP97088. 10.7554/eLife.97088.2

Miller, C. T., Freiwald, W. A., Leopold, D. A., Mitchell, J. F., Silva, A. C., & Wang, X. (2016). Marmosets: A Neuroscientific Model of Human Social Behavior. Neuron, 90(2), 219–233. doi:10.1016/j.neuron.2016.03.018

Miller, C. T., Gire, D., Hoke, K., Huk, A. C., Kelley, D., Leopold, D. A., … Niell, C. M. (2022). Natural behavior is the language of the brain. Curr Biol, 32(10), R482–r493. doi:10.1016/j.cub.2022.03.031

Miss, F. M., & Burkart, J. M. (2018). Corepresentation during joint action in marmoset monkeys (Callithrix jacchus). Psychological Science, 29(6), 984–995.

Mitchell, J. F., Reynolds, J. H., & Miller, C. T. (2014). Active vision in marmosets: a model system for visual neuroscience. J Neurosci, 34(4), 1183–1194. doi:10.1523/jneurosci.3899-13.2014

Mosher, C. P., Zimmerman, P. E., & Gothard, K. M. (2014). Neurons in the monkey amygdala detect eye contact during naturalistic social interactions. Curr Biol, 24(20), 2459–2464. doi:10.1016/j.cub.2014.08.063

Mustoe, A. (2023). A tale of two hierarchies: Hormonal and behavioral factors underlying sex differences in social dominance in cooperative breeding callitrichids. Horm Behav, 147, 105293. doi:10.1016/j.yhbeh.2022.105293

Ngo, V., Gorman, J. C., De la Fuente, M. F., Souto, A., Schiel, N., & Miller, C. T. (2022). Active vision during prey capture in wild marmoset monkeys. Curr Biol, 32(15), 3423–3428.e3423. doi:10.1016/j.cub.2022.06.028

Pandey, S., Simhadri, S., & Zhou, Y. (2020). Rapid Head Movements in Common Marmoset Monkeys. iScience, 23(2), 100837. doi:10.1016/j.isci.2020.100837

Pereira, T. D., Tabris, N., Matsliah, A., Turner, D. M., Li, J., Ravindranath, S., … Murthy, M. (2022). SLEAP: A deep learning system for multi-animal pose tracking. Nature Methods, 19(4), 486–495. doi:10.1038/s41592-022-01426-1

Pfister, R., Schwarz, K. A., Janczyk, M., Dale, R., & Freeman, J. (2013). Good things peak in pairs: a note on the bimodality coefficient. Frontiers in Psychology, 4. doi:10.3389/fpsyg.2013.00700

Ramezanpour, H., & Thier, P. (2020). Decoding of the other’s focus of attention by a temporal cortex module. Proc Natl Acad Sci U S A, 117(5), 2663–2670. doi:10.1073/pnas.1911269117

Saxe, R., & Kanwisher, N. (2003). People thinking about thinking people. The role of the temporo-parietal junction in “theory of mind”. NeuroImage, 19(4), 1835–1842. doi:10.1016/s1053-8119(03)00230-1

Shepherd, S. (2010). Following Gaze: Gaze-Following Behavior as a Window into Social Cognition. Frontiers in Integrative Neuroscience, 4. doi:10.3389/fnint.2010.00005

Shepherd, S. V., Deaner, R. O., & Platt, M. L. (2006). Social status gates social attention in monkeys. Curr Biol, 16(4), R119–120. doi:10.1016/j.cub.2006.02.013

Shepherd, S. V., & Freiwald, W. A. (2018). Functional Networks for Social Communication in the Macaque Monkey. Neuron, 99(2), 413–420.e413. doi:10.1016/j.neuron.2018.06.027

Solomon, N. G., & French, J. A. (1997). *Cooperative breeding in mammals*: Cambridge University Press.

Sturman, O., von Ziegler, L., Schläppi, C., Akyol, F., Privitera, M., Slominski, D., … & Bohacek, J. (2020). Deep learning-based behavioral analysis reaches human accuracy and is capable of outperforming commercial solutions. Neuropsychopharmacology, 45(11), 1942–1952.

Spadacenta, S., Dicke, P.W. & Thier, P. (2019). Reflexive gaze following in common marmoset monkeys. Sci Rep 9, 15292. 10.1038/s41598-019-51783-9

Spadacenta, S., Dicke, P. W., & Thier, P. (2022). A prosocial function of head-gaze aversion and head-cocking in common marmosets. Primates, 63(5), 535–546. doi:10.1007/s10329-022-00997-z

Tomasello, M., Carpenter, M., Call, J., Behne, T., & Moll, H. (2005). Understanding and sharing intentions: the origins of cultural cognition. The Behavioral and brain sciences, 28(5), 675–735. 10.1017/S0140525X05000129

Yamamoto, M. E., Araujo, A., Arruda, M. d. F., Lima, A. K. M., Siqueira, J. d. O., & Hattori, W. T. (2014). Male and female breeding strategies in a cooperative primate. Behavioural Processes, 109, 27–33. 10.1016/j.beproc.2014.06.009

